# MAST-Decon: Smooth Cell-type Deconvolution Method for Spatial Transcriptomics Data

**DOI:** 10.1101/2024.05.10.593595

**Authors:** Tianyou Luo, Jiawen Chen, Wenrong Wu, Jinying Zhao, Huaxiu Yao, Hongtu Zhu, Yun Li

## Abstract

Spatial transcriptomics (ST) technologies have gained increasing popularity due to their ability to provide positional context of gene expressions in a tissue. One major limitation of current commercially available ST methods such as the 10X Genomics Visium platform is the lack of single cell resolution. Cell type deconvolution for ST data is critical in order to fully reveal underlying biological mechanisms. Existing ST data deconvolution methods share two common limitations: first, few of them utilize spatial neighborhood information. Existing methods such as RCTD and SPOTlight intrinsically treat each spatial spot as independent of neighboring spots, although we anticipate nearby spots to share similar cell type compositions based on clinical evidence of tissue structures. Such limitation could be amplified when sequencing depths at single spots are relatively low so that borrowing information from neighboring spots is necessary in order to obtain reliable deconvolution results. Second, although Visium data provide us with a histological image which could add additional information regarding spot heterogeneity, most existing methods do not utilize this H&E image. To solve these two limitations, we developed Multiscale Adaptive ST Deconvolution (MAST-Decon), a smooth deconvolution method for ST data. MAST-Decon uses a weighted likelihood approach and incorporates both gene expression data, spatial neighborhood information and H&E image features by constructing different kernel functions to obtain a smooth deconvolution result. We showcased the strength of MAST-Decon through simulations based on real data, including a single-cell dataset of mouse brain primary visual cortex, and real-world Visium datasets to demonstrate its robust and superior performance compared with other state-of-the-art methods.

## 1 Introduction

Measurement of gene expression levels plays an important and central role in the characterization of cells and their surrounding tissues. Next-generation sequencing (NGS) of bulk cell populations and single cells has become an ubiquitous tool in modern-day biological research, with numerous contributions in advancing our understandings of many biological mechanisms and functions. However, these technologies require dissolving the cells first, leading to the loss of information about the spatial structure, interaction and spatial relationships between the cells. In recent years, spatial transcriptomics (ST) technologies have gained increasing popularity due to their ability to provide positional context of gene expressions in the tissue. Retaining spatial context is especially important in exploring spatially heterogeneous structures such as brain cortex and tumor microenvironments.

Current ST technologies largely fall into two categories, namely imaging-based approaches and NGS-based technologies [1, 2]. Imaging-based technologies include methods based on fluorescence i n s itu hybridization ( FISH) s uch as multiplexed error-robust FISH (MERFISH) [3, 4], sequential FISH (seqFISH) [5, 6] and seqFISH+ [7], which obtain quantitative gene expression counts by directly imaging individual RNA molecules. They also include in situ sequencing (ISS)-based methods [8] such as STARmap [9], in which RNAs were reverse transcribed, amplified and sequenced by ligation in situ. Imaging-based technologies can reach cellular or even subcellular resolution, but due to the limitations arising from the imaging step, they typically require gene selection in advance. Despite several efforts to scale up to about 10,000 genes [4, 7], most of the time they can only target hundreds of genes in practice. The complexity of these experiments and the amount of effort needed to optimize the experimental workflow have limited the more expansive usage of such methods. NGS-based methods, such as the Spatial Transcriptomics platform [10] and Visium platform by 10X Genomics and Slide-seq [11, 12], rely on spatial barcoding. Commercially available NGS-based methods are widely accessible to researchers and can target the whole transcriptome without prior gene selection, but they cannot yet reach single-cell resolution. The number of cells within a spatial spot may range from 1 to 200 depending on the biological tissue and ST platform. Therefore, any downstream analysis such as spatially variable gene detection could be confounded by differential cell type compositions across spots. To solve this problem of current NGS-based ST technologies, many cell type deconvolution methods have been developed in order to fully reveal underlying biological structures. These methods utilize an external single-cell or single-nucleus RNA-sequencing (scRNA-seq or snRNA-seq) reference dataset to infer the cell-type composition for each spot.

Existing ST data deconvolution methods share two common limitations: first, few of them utilize spatial neighborhood information. Existing methods such as RCTD and SPOTlight intrinsically treat each spatial spot as independent of neighboring spots, although we anticipate nearby spots to share similar cell type compositions based on clinical evidence of tissue structures. Such limitation could be amplified when sequencing depths at single spots are relatively low so that borrowing information from neighboring spots is necessary in order to obtain reliable deconvolution results. Second, although Visium data provide us with a histological image which could add additional information regarding spot heterogeneity, most existing methods do not utilize this hematoxylin and eosin (H&E) stained image. To solve these two limitations, we developed Multiscale Adaptive ST Deconvolution (MAST-Decon), a smooth deconvolution method for ST data. MAST-Decon uses a weighted likelihood approach and incorporates both gene expression data, spatial neighborhood information and H&E image features by constructing different kernel functions to obtain a smooth deconvolution result. We showcased the strength of MAST-Decon through a simulated benchmark dataset. By introducing spatial smoothness, we were able to correct several spots erroneously inferred by RCTD and achieved better deconvolution results than state-of-the-art deconvolution methods as measured by root mean-squared error (RMSE) and Spearman correlation in simulation. We also applied MAST-Decon to real data from mouse somatosensory cortex, human brain dorsolateral prefrontal cortex (DLPFC) and hepatocellular carcinoma (HCC). Our results show that MAST-Decon provides robust cell type proportion estimates across various data qualities and tissue types.

## 2 Results

### 2.1 Smooth cell-type deconvolution of Spatial Transcriptomics data using MAST-Decon

MAST-Decon is motivated by the Multiscale Adaptive Regression Model (MARM) [13] in medical imaging analysis. We aim to perform smooth cell-type deconvolution for ST data by flexibly borrowing information from neighboring spots. Specifically, the first step is to specify the probabilistic model for each individual spot, where we largely follow the Poisson model used in RCTD. For each spot *d* ∈ **D** and gene *i* ∈ **G**, where **D** and **G** are the set of all spots and all genes respectively, the observed gene counts *Y*_*i*_(*d*) are assumed to follow a Poisson distribution depending on the spot’s total UMI count *N*_*d*_ and the rate parameter *λ*_*i*_(*d*) which characterizes the mixture of different cell types:

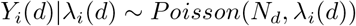

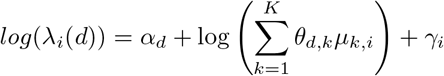

Here *µ*_*k,i*_ is the mean gene expression profile for the *k*th cell-type inferred from external single-cell reference. *γ*_*i*_ is a gene-specific random effect introduced by RCTD accounting for platform differences between ST data and reference scRNA-seq/snRNA-seq data. Then the log-likelihood for spot *d* can be written as:

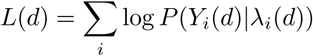

Existing methods utilize various optimization methods to obtain the maximum likelihood estimator of *θ*_*d,k*_ to make inference about the cell-type proportions for each spot. Notice that in this formulation of the log-likelihood, each spot only involves its own gene expression data and parameters, and the estimation is carried out for each spot separately without any input from neighboring spots. Instead of only considering the target spot’s gene expression, it could be more efficient to utilize all information in the spatial neighborhood of each target spot. Taking advantage of this, MAST-Decon chooses to optimize a weighted log-likelihood for each spot:

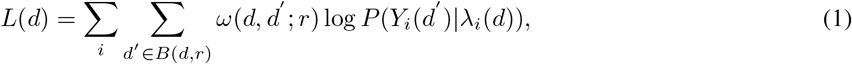

where the weight function *ω*(*d, d*′ ; *r*) characterizes the similarities between two spots *d* and *d*′, and *r* is the radius to determine how many neighboring spots should be used in calculating the weighted log-likelihood. We factorize the weight function into three different parts:

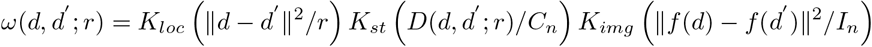

*K*_*loc*_(*u*), *K*_*st*_(*u*) and *K*_*img*_(*u*) are three non-negative kernel functions with compact support such that all of them decrease to 0 as *u* increases. The first term represents the similarity between two spots in terms of spatial distance. Nearby spots are expected to be more similar in terms of cell-type compositions than spots further away from each other, therefore this kernel function gives weights inversely proportional to the distance between spot *d* and *d*^*′*^. In practice, we set the kernel function to be

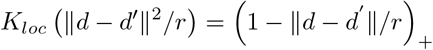

Therefore, we only consider spots within a ball of the current specified radius, and far away spots will not be included to avoid over-smoothing the pattern. The second term downweights the spots with large *D*_*θ*_(*d, d*′ ; *r*) values. In practice, we use the following equation for this distance metric:

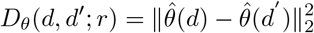

This kernel measures the difference between current cell type proportion estimates 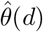 and 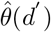. For any given spot of interest, any other spots with very different inferred cell type proportions will have little impact on its likelihood function. In practice, the kernel is set to *K*_*st*_(*u*) = exp(*−u*). By doing so, neighboring spots with highly different cell type proportions (such as spots on the boundary of different cortical layers) will not be over-smoothed, and a clear tissue boundary can be retained. Parameter *C*_*n*_ controls the smoothness of inferred results. In practice we set it to be a tuning parameter *C* times the median of *D*_*θ*_(*d, d*′ ; *r*) in the ST dataset. A larger *C* will dilute the effect of *D*_*θ*_(*d, d*′ ; *r*) and tend to create a uniformly large weight for neighboring spots, encouraging higher smoothness among the results. A smaller *C* instead will dilate the effect of spot similarity distance, creating large weights only for highly similar spots. The choice of the tuning parameter is discussed in our simulation studies.

The third kernel is optional and measures how much the spots differ in terms of other information such as histological imaging features, based on the embedded imaging feature vectors of each spot which can be obtained using some existing pretrained deep learning models. Note that the weight calculation is flexible and adaptive, meaning that it can be modified according to the information we have as well. If we do not have reliable histological information, the third kernel can be eliminated and only the location kernel and ST kernel are used for calculating the weights. Furthermore, if we have clinical annotations that can be used to guide the deconvolution, we could also use that as an input matrix instead of the image kernel.

The inference procedure of MAST-Decon is carried out in an iterative way: we first specify the series of radii to be used. Typically this can be set to the distances in the ST dataset between nearest spots, second nearest spots and so on. We start with an initial deconvolution result with *r* = 0, which corresponds to not incorporating any neighborhood information. This initial result can be easily obtained using RCTD algorithm and the estimated parameters 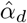 and 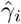 will be fixed afterwards. Then we calculate the weight functions based on the current estimates of cell type proportions 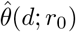 and the next radius in the series *r*_1_, and construct weighted log-likelihood for each spot. The weighted log-likelihood for each spot is maximized to obtain a new set of cell type proportion estimates, which is again combined with the next radius in the series to calculate the updated weights and weighted log-likelihoods. This procedure is iterated until we reach the last radius in the pre-specified series. This process is summarized in Figure 1 and Algorithm 1.

**Figure 1:**
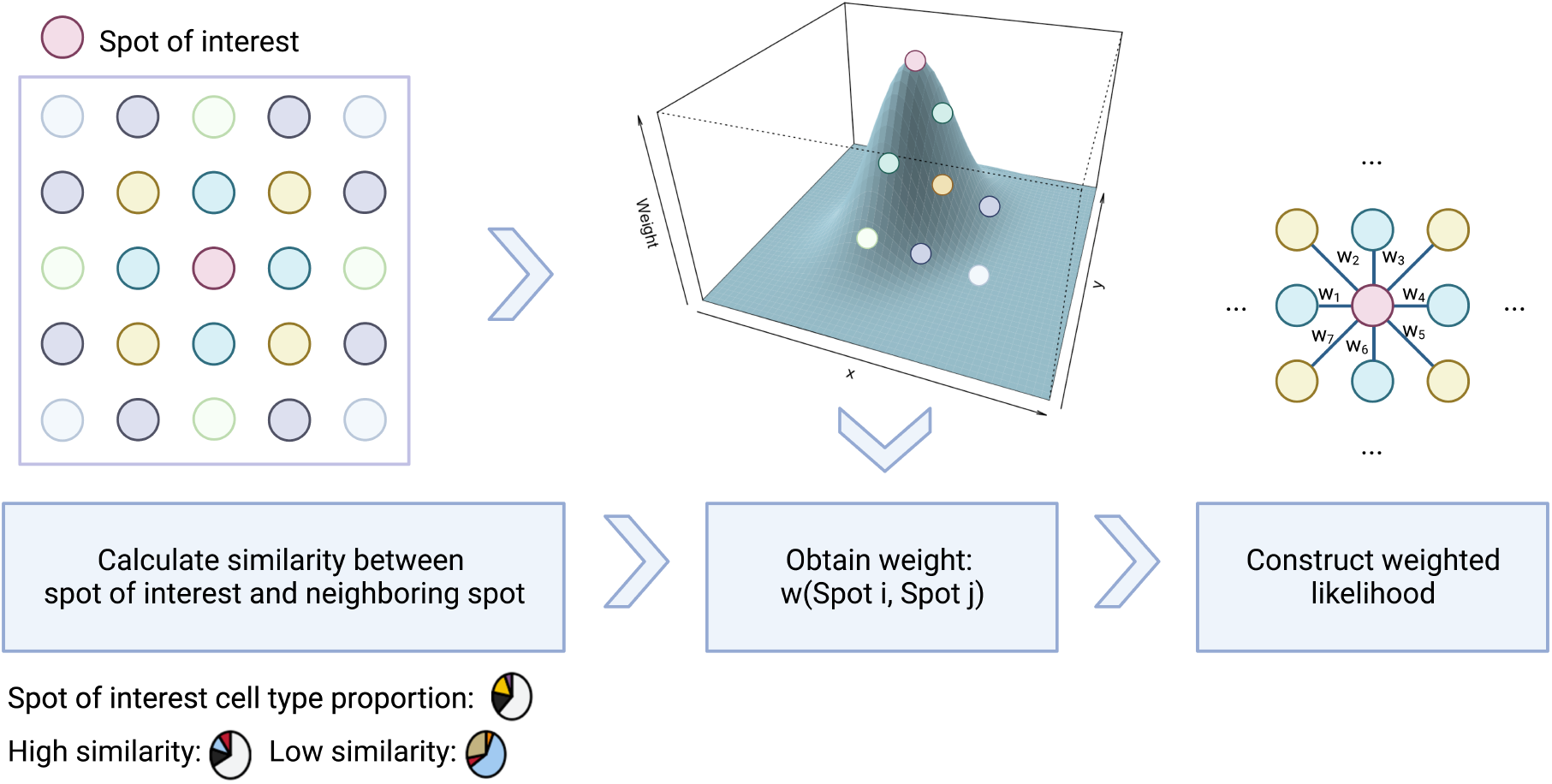
Schematic representation of MAST-Decon workflow. After each iteration, the weights for each neighboring spot are re-calculated based on the distances and cell type proportion similarities between the two spots, and new neighboring spots are included based on a new radius (blue→yellow→green→purple spots in the left figure). The weighted likelihoods for each spot are updated using newly calculated weights, and optimized in the next iteration to obtain updated cell type proportion estimates.

#### Algorithm 1: MAST-Decon inference algorithm

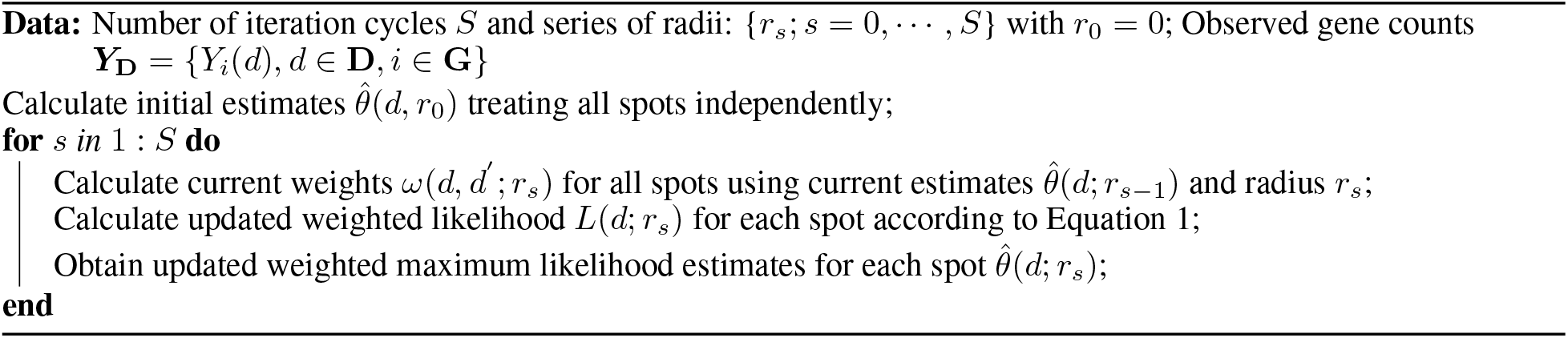

### 2.2 MAST-Decon outperforms state-of-the-art methods in simulated benchmark data

We first conducted a proof-of-concept simulation to demonstrate the advantage of MAST-Decon over existing ST deconvolution methods. Following discoveries from previous benchmarking studies [14, 15], we included our baseline method RCTD [16], two other state-of-the-art deconvolution methods stereoscope [17] and Cell2location [18], and another spatially-aware ST deconvolution method CARD [19] in our comparison. We created a simulated benchmark dataset using single-cell RNA sequencing (scRNA-seq) data from the mouse brain primary visual cortex [20]. The scRNA-seq data were randomly split into two equal subsets. One subset was used to create pseudo ST spots and the other half served as the scRNA-seq reference in deconvolution. This allowed for an ideal scenario where matched references are available and no batch effects are present. Since the scRNA-seq data were generated following the micro-dissection of cells into different cortical layers, we localized the cells according to their respective layers and created pseudo ST spots by randomly selecting cells from the corresponding cortical layers, pooling the gene counts together and randomly downsampling to a predetermined sequencing depth, akin to that observed in real Visium data (a mean of 2,500 UMI counts). We used the spot-level root mean squared error (RMSE) and Spearman correlation between the estimated and ground-truth cell-type proportions as evaluation metrics.

While all methods correctly inferred the layered structure of cell type distributions, we observed multiple spots whose RCTD-inferred cell type proportions did not align with the expected pattern in the corresponding layer. Conversely, MAST-Decon consistently produced cell type proportion estimates that closely matched the ground truth (as highlighted in Figure 2a-c). Cell2location achieved similar or better performances than RCTD, but is still outperformed by MAST-Decon. Stereoscope underestimated the proportion of L4 cells in many Layer 4 spots. CARD tended to underestimate the proportions of cell types specific to their corresponding layers while overestimate the proportions of cell types localized in the neighboring layers. This might be attributed to slight over-smoothing resulting from its parametric spatial dependence model. Across all cortical layers, MAST-Decon achieved the lowest RMSE and highest Spearman Correlation (Figure 2d-e).

**Figure 2:**
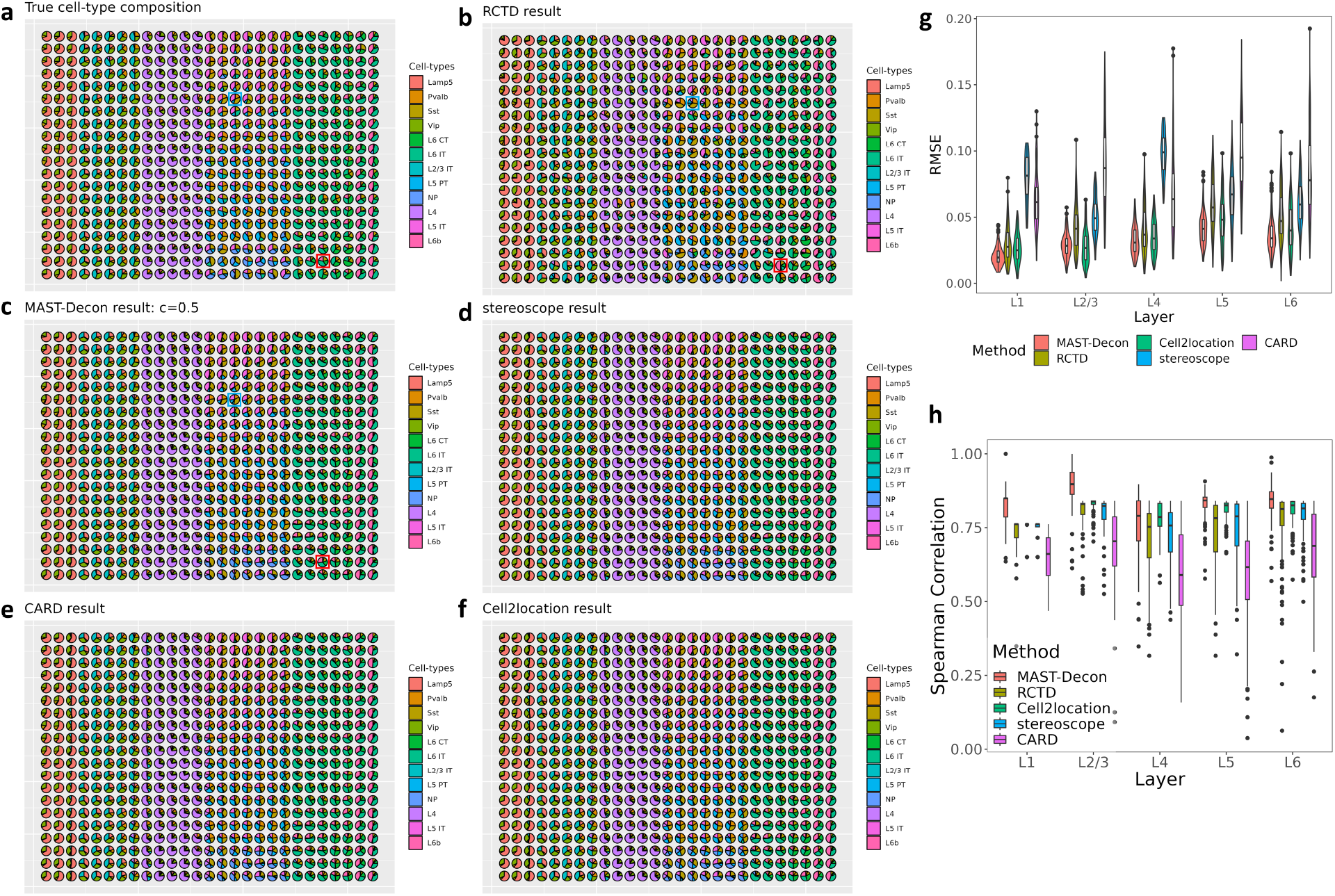
Comparing MAST-Decon with existing methods on a dataset simulated using mouse brain scRNA-seq data. **(a)** True cell type composition of each simulated spot. **(b)-(f)** Cell type compositions inferred by **(b)** RCTD, **(c)** MAST-Decon, **(d)** stereoscope, **(e)** CARD and **(f)** Cell2location. Blue and red squares in **(a)-(c)** highlight example spots where RCTD inferred cell type proportions do not match the ground truth nor expected patterns of the corresponding layer, while MAST-Decon inferred cell-type compositions align well with the ground truth. **(g)** RMSE and **(h)** Spearman correlation between inferred and ground truth cell type proportions for each spot across different layers.

### 2.3 MAST-Decon robustly uncovers cell-type structure with down-sampled and real mouse and human brain Visium data

One potential advantage of MAST-Decon over existing methods is its ability to efficiently handle ST data with low sequencing depth by incorporating information from neighboring spots. To demonstrate such an advantage of MAST-Decon, we down-sampled Visium ST data from mouse somatosensory cortex [18] to different sequencing depths and evaluated the performances of various methods under different data sparsity levels. For each slide, we manually annotated the spots to different cortical layers according to histological images and marker gene expressions. Since the ground truth cell type proportions of each spot are not available, for evaluation purposes, we performed *k*-means clustering on inferred cell type proportions to assign each spot to a cluster and computed Adjusted Rand Index (ARI) against the manual annotation. A higher ARI indicates a stronger agreement between deconvolution results and the expected cortical layer structure. As shown in Figure 3, MAST-Decon significantly outperformed existing methods and retained a robust deconvolution performance across different sequencing depths, and revealed much clearer cortical layer structure as expected. MAST-Decon with three different smoothing levels performed similarly when the mean UMI count is not too low. For the lowest sequencing depth (mean UMI count = 1000) scenario, MAST-Decon with higher smoothing levels performs better than MAST-Decon with lower smoothing (*C* = 0.5). This is expected since with very shallow sequencing depth, a higher smoothing level is necessary to borrow enough information from neighboring spots in order to uncover the underlying structure. RCTD and stereoscope perform reasonably well when the data are of high sequencing depth and mean UMI counts per spot are high, but their performance deteriorates when the ST data are of a relatively lower sequencing depth. CARD appears to have over-smoothed the cell type composition patterns, with the proportion of L2/3 cells gradually decreasing and the proportion of L6 cells gradually increasing from the outer layer to inner layer, but mixing up the layer structures in the middle. Cell2location fails to produce a reasonable set of results in our setting, potentially suggesting sensitivity to the choice of scRNA-seq reference data.

**Figure 3:**
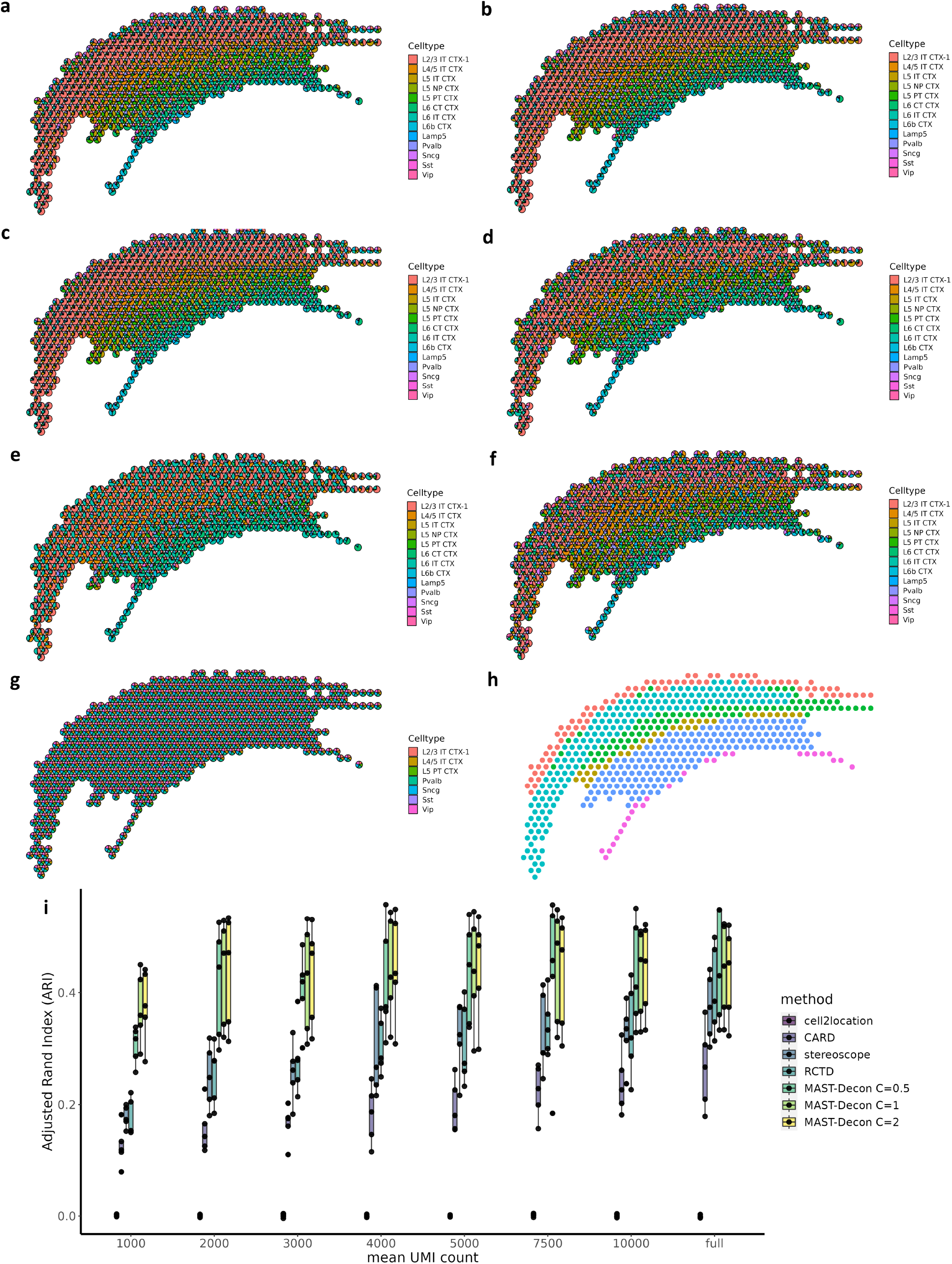
Comparing MAST-Decon with existing deconvolution methods on a real mouse brain Visium dataset. **(a)-(g)**: Cell-type composition inferred for one slide (ST8059048) downsampled to a mean UMI count of 2,000 by **(a)** MAST-Decon with *C* = 0.5 **(b)** MAST-Decon with *C* = 1 **(c)** MAST-Decon with *C* = 2 **(d)** RCTD **(e)** CARD **(f)** stereoscope **(g)** cell2location; **(i)** Adjusted Rand Index (ARI) for different methods across different sequencing depths. ARIs are computed against pathologist’s layer annotations **(h)** using *k*-means clustering results from inferred cell type proportions.

In addition to downsampling high-depth data, we also evaluated our method using real Visium ST data from postmortem human dorsolateral prefrontal cortex (DLPFC) [21, 22]. The dataset includes twelve slides in six pairs of two ‘spatial replicates’ from three adult donors. We utilized snRNA-seq data from human brain DLPFC [23] as reference, consisting of 54,151 cells from 13 cell types. The spots were manually annotated to six different cortical layers (L1-L6) and white matter (WM), enabling us to investigate the distributions of inferred cell type proportions in different cortical regions. ARIs computed against the layer annotations across twelve slides show that MAST-Decon results display clearer cortical structures compared to RCTD and CARD (Figure 4a). Here we focus on slide 151674 (median UMI per spot = 5,296) (Figure 4b-c) with results for additional slides available in the supplementary files. As shown in stacked violin plots and cell type proportion heatmaps (Figure 4d-e), MAST-Decon inferred cell type proportions of layer-specific cell types are more enriched in the corresponding cortical layers compared to RCTD’s estimations. Notably, RCTD estimates show L4 excitatory neurons (Excit_L4) spread across layer 2 to layer 5, whereas MAST-Decon results localize them mainly in layer 4 and layer 5. Proportions of L3, L5 and L6 excitatory neurons estimated by MAST-Decon exhibit clearer enrichment in their respective layers as well. Astrocytes (Astro) are enriched in layer 1, while oligodendrocytes (Oligo) are predominantly found in white matter regions. CARD results (Figure 4f) again show clear over-smoothing effects. Oligodendrocytes are inferred to be present across all cortical layers instead of being localized in white matter, and estimated astrocyte proportions are much higher than expected in L2-L6. Across different cortical regions, the spatial distributions of MAST-Decon inferred cell type proportions appear more reasonable and align better with manual annotations and expected cortical patterns.

**Figure 4:**
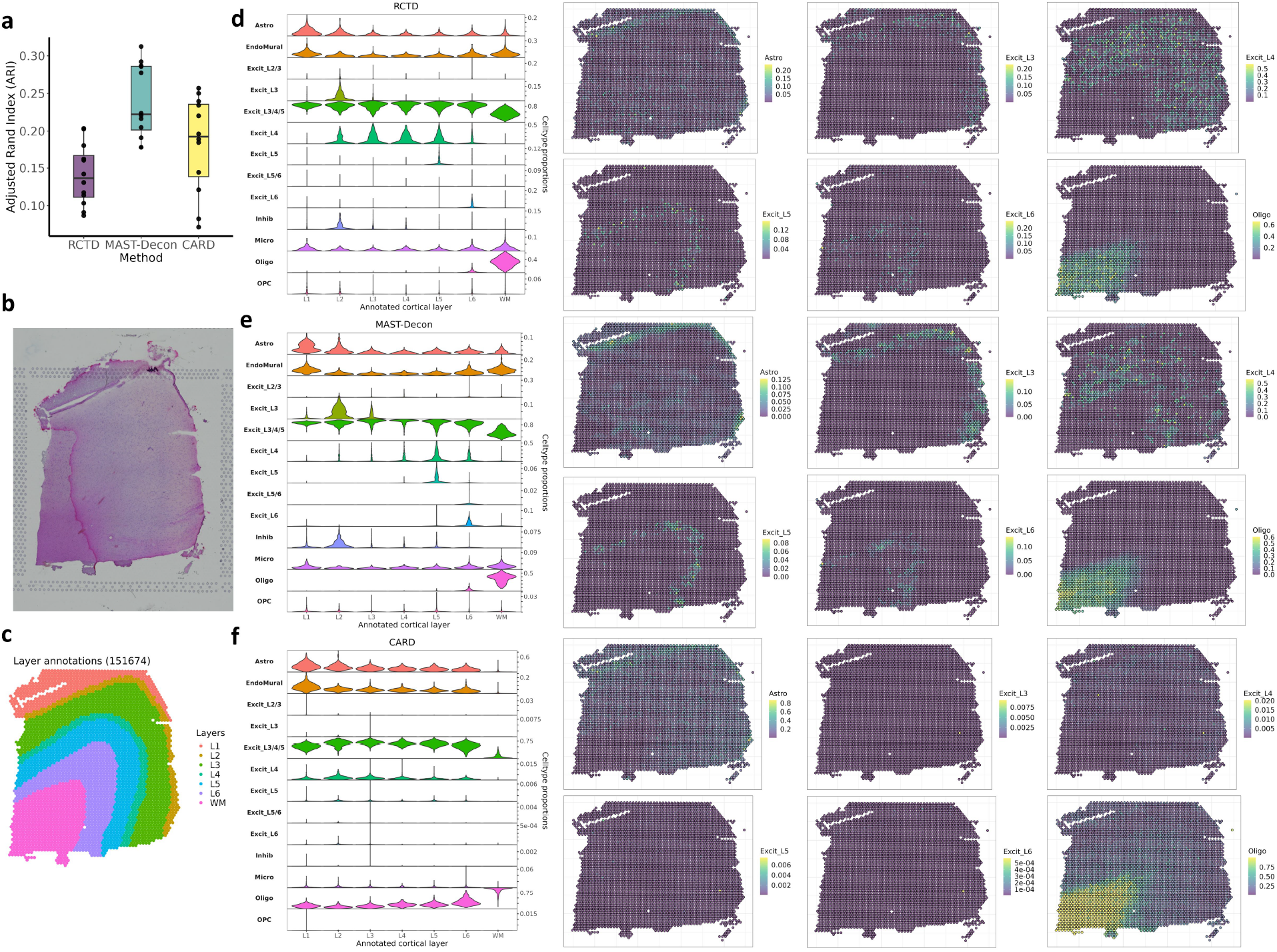
Comparing MAST-Decon with RCTD and CARD on human postmortem dorsolateral prefrontal cortex (DLPFC) data. **(a)** Adjusted Rand Index (ARI) for different methods computed on 12 slides. ARIs were computed against manual layer annotations using *k*-means clustering results from inferred cell type proportions. **(b)** Low-resolution histology image of slide 151674. **(c)** Layer annotations of slide 151674. **(d)-(f)**: Cell type proportions for slide 151674 inferred by **(d)** RCTD, **(e)** MAST-Decon, and **(f)** CARD. Stacked violin plots on the left show the distributions of estimated proportions of each cell type in each of the cortical layers (L1-L6) and white matter (WM). Heatmaps on the right display the estimated spatial distributions of six selected layer-specific cell types.

### 2.4 MAST-Decon identifies tertiary lymphoid structure in human hepatocellular carcinoma

We further applied MAST-Decon to analyze ST data obtained from a tissue section of a hepatocellular carcinoma (HCC) patient [24]. The tumor nodule from patient HCC-5 in the original dataset was cut into four parts for Visium sequencing, and we focus on section HCC-5C in our analysis. We used the HCCDB-SC3 dataset from HCCDB v2.0 [25] as scRNA-seq reference for deconvolution, which was obtained from seven tumor samples and three tumor-adjacent liver tissues in seven HCC patients. The scRNA-seq reference data consist of 61,370 cells from 16 cell types. The spots were annotated into four different tissue regions, including immune, tumor, normal and stromal (Figure 5a). ARI computed against region annotations was much higher for MAST-Decon results (ARI=0.575) compared to RCTD results (ARI=0.228), indicating MAST-Decon’s superior ability to dissect different tissue regions. Tertiary lymphoid structures (TLS) are lymphoid aggregates that form in response to inflammatory processes. They play a crucial role in anti-tumor immunotherapeutic responses and have been associated with improved clinical outcomes [26]. MAST-Decon accurately identified the TLS in the tissue section. The deconvolution results revealed that the TLS spots consist of a high proportion of B cells, T cells (CD8+ T cells, Tregs and circulating NK/NKT cells) and cDCs (conventional dendritic cells), which is consistent with the enrichment analysis findings in the original paper (Figure 5b-c). We also observed the existence of many immune cells such as CD4+ T cells, CD8+ T cells, cDCs, circulating NK/NK-T cells, monocytes & monocyte-derived cells and Tregs in the stromal region. Notably, MAST-Decon estimated higher cell type proportions of CD8+ T cells, cDCs, Tregs and monocytes in the stromal region than those estimated by RCTD (Figure 6). Although we do not know the ground truth cell type distributions of this slide, a close examination of some of the immune cell markers suggests a rich presentation of immune cells in the stromal region (Figure 5d). The existence of immune cell types such as monocytes in peritumoral stroma of HCC can create a distinct tumor microenvironment and play an important role in therapeutic response and disease progression [27, 28]. We also note that the annotated normal spots are estimated to have a high proportion of malignant cells as well. This is potentially due to the fact that all scRNA-seq reference data come from HCC patients and therefore normal liver cells are under-represented in the reference.

**Figure 5:**
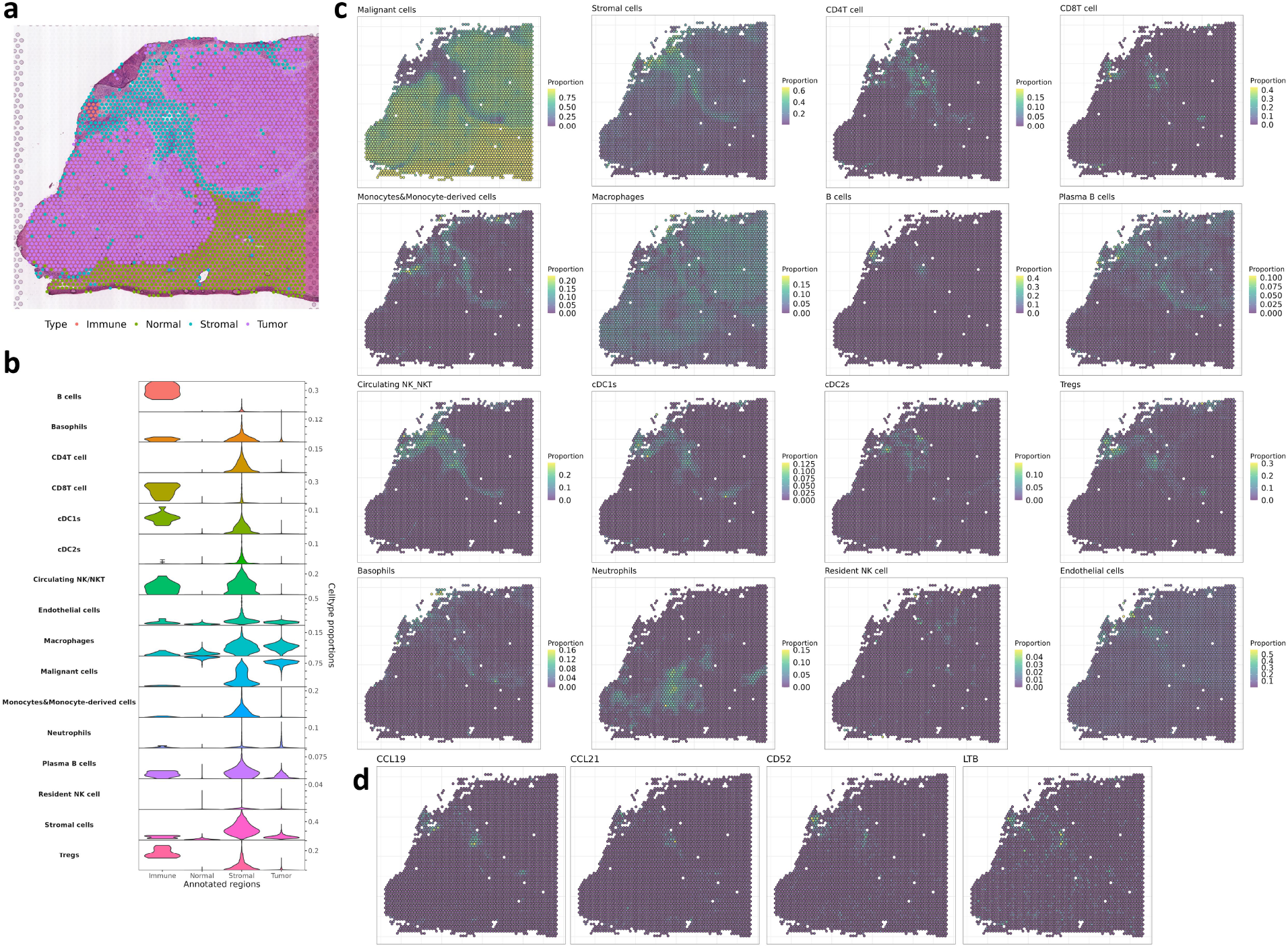
Applying MAST-Decon to human hepatocellular carcinoma (HCC) Visium data. **(a)** Manual regional annotations overlaid on the histology image of slide HCC-5C. **(b)** Stacked violin plot showing the distributions of proportions of each cell type in each of the four regions as estimated by MAST-Decon. **(c)** Heatmaps displaying the spatial distributions of different cell types estimated by MAST-Decon. **(d)** Expression levels of selected genes.

**Figure 6:**
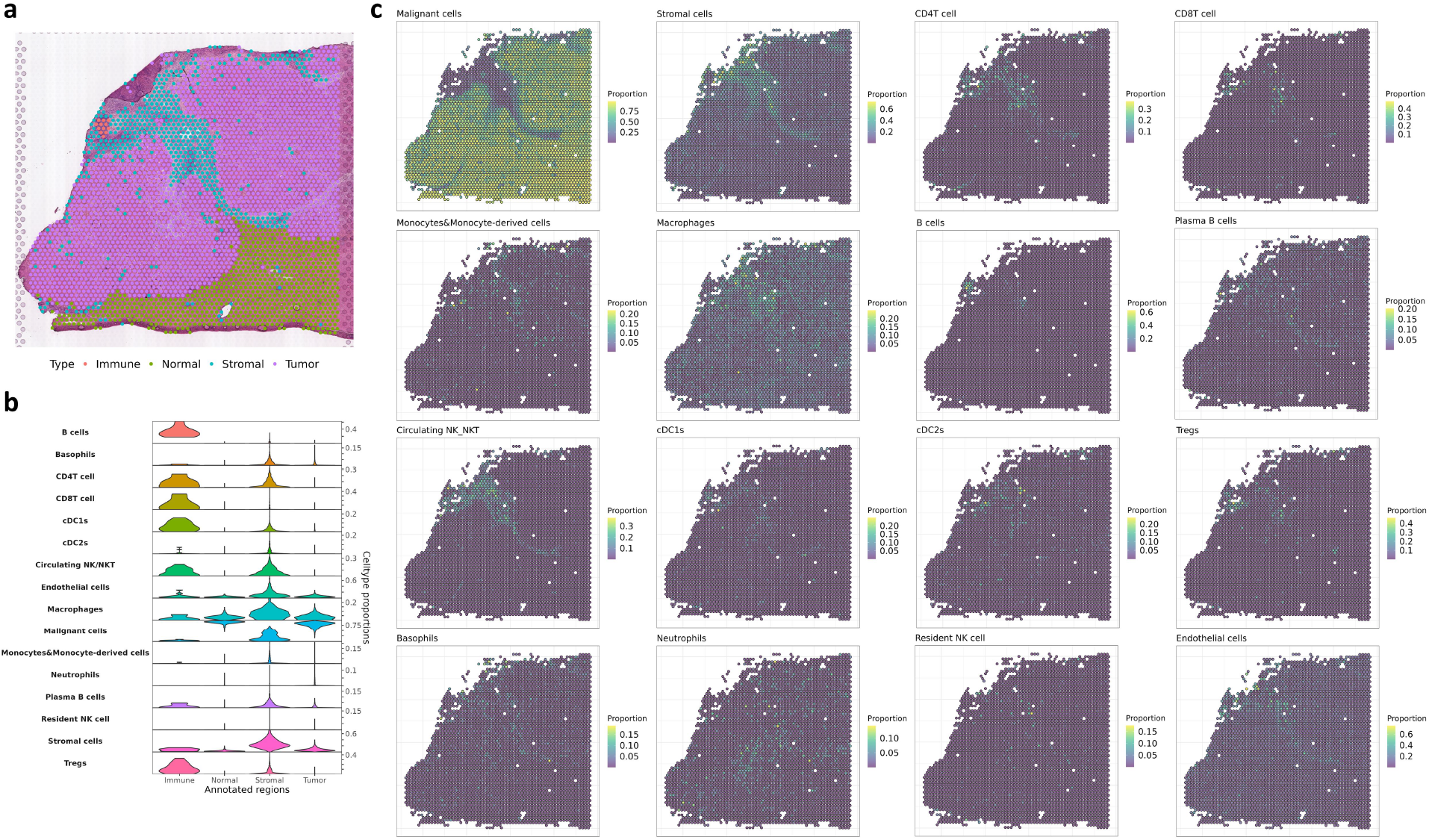
Applying RCTD to human hepatocellular carcinoma (HCC) Visium data. **(a)** Manual regional annotations overlaid on the histology image of slide HCC-5C. **(b)** Stacked violin plot showing the distributions of proportions of each cell type in each of the four regions as estimated by RCTD. **(c)** Heatmaps displaying the spatial distributions of different cell types estimated by RCTD.

## 3 Discussion

Existing spatial transcriptomics data deconvolution methods typically treat the spots separately and therefore ignore the spatial neighborhood information and spatial correlation in cell-types between neighboring spots. Here we introduce MAST-Decon, a smooth cell-type deconvolution method for ST data which utilizes expression profiles and spatial neighborhood information together to achieve better deconvolution results. MAST-Decon integrates information from neighboring spots through an adaptive weighted likelihood approach, therefore avoiding explicitly specifying spatial dependency structure. Through simulations and real data analysis, we show that MAST-Decon achieves more reliable and robust deconvolution results compared to existing deconvolution methods.

## References

[1] Rao, A., Barkley, D., França, G. S., & Yanai, I. (2021). Exploring tissue architecture using spatial transcriptomics. Nature, 596(7871), 211–220.

[2] Moses, L., & Pachter, L. (2022). Museum of spatial transcriptomics. Nature Methods, 19(5), 534–546.

[3] Chen, K. H., Boettiger, A. N., Moffitt, J. R., Wang, S., & Zhuang, X. (2015). Spatially resolved, highly multiplexed rna profiling in single cells. Science, 348(6233), aaa6090.

[4] Xia, C., Fan, J., Emanuel, G., Hao, J., & Zhuang, X. (2019). Spatial transcriptome profiling by merfish reveals subcellular rna compartmentalization and cell cycle-dependent gene expression. Proceedings of the National Academy of Sciences, 116(39), 19490–19499.

[5] Lubeck, E., Coskun, A. F., Zhiyentayev, T., Ahmad, M., & Cai, L. (2014). Single-cell in situ RNA profiling by sequential hybridization. Nature methods, 11(4), 360–361.

[6] Shah, S., Lubeck, E., Zhou, W., & Cai, L. (2016). In situ transcription profiling of single cells reveals spatial organization of cells in the mouse hippocampus. Neuron, 92(2), 342–357.

[7] Eng, C.-H. L., Lawson, M., Zhu, Q., Dries, R., Koulena, N., Takei, Y., Yun, J., Cronin, C., Karp, C., Yuan, G.-C., et al. (2019). Transcriptome-scale super-resolved imaging in tissues by RNA seqFISH+. Nature, 568(7751), 235–239.

[8] Ke, R., Mignardi, M., Pacureanu, A., Svedlund, J., Botling, J., Wählby, C., & Nilsson, M. (2013). In situ sequencing for RNA analysis in preserved tissue and cells. Nature methods, 10(9), 857–860.

[9] Wang, X., Allen, W. E., Wright, M. A., Sylwestrak, E. L., Samusik, N., Vesuna, S., Evans, K., Liu, C., Ramakrishnan, C., Liu, J., et al. (2018). Three-dimensional intact-tissue sequencing of single-cell transcriptional states. Science, 361(6400), eaat5691.

[10] Ståhl, P. L., Salmén, F., Vickovic, S., Lundmark, A., Navarro, J. F., Magnusson, J., Giacomello, S., Asp, M., Westholm, J. O., Huss, M., et al. (2016). Visualization and analysis of gene expression in tissue sections by spatial transcriptomics. Science, 353(6294), 78–82.

[11] Rodriques, S. G., Stickels, R. R., Goeva, A., Martin, C. A., Murray, E., Vanderburg, C. R., Welch, J., Chen, L. M., Chen, F., & Macosko, E. Z. (2019). Slide-seq: A scalable technology for measuring genome-wide expression at high spatial resolution. Science, 363(6434), 1463–1467.

[12] Stickels, R. R., Murray, E., Kumar, P., Li, J., Marshall, J. L., Di Bella, D. J., Arlotta, P., Macosko, E. Z., & Chen, F. (2021). Highly sensitive spatial transcriptomics at near-cellular resolution with Slide-seqV2. Nature biotechnology, 39(3), 313–319.

[13] Li, Y., Zhu, H., Shen, D., Lin, W., Gilmore, J. H., & Ibrahim, J. G. (2011). Multiscale adaptive regression models for neuroimaging data. Journal of the Royal Statistical Society: Series B (Statistical Methodology), 73(4), 559–578.

[14] Chen, J., Liu, W., Luo, T., Yu, Z., Jiang, M., Wen, J., Gupta, G. P., Giusti, P., Zhu, H., Yang, Y., et al. (2022). A comprehensive comparison on cell-type composition inference for spatial transcriptomics data. Briefings in Bioinformatics, 23(4), bbac245.

[15] Li, B., Zhang, W., Guo, C., Xu, H., Li, L., Fang, M., Hu, Y., Zhang, X., Yao, X., Tang, M., et al. (2022). Benchmarking spatial and single-cell transcriptomics integration methods for transcript distribution prediction and cell type deconvolution. Nature Methods, 1–9.

[16] Cable, D. M., Murray, E., Zou, L. S., Goeva, A., Macosko, E. Z., Chen, F., & Irizarry, R. A. (2022). Robust decomposition of cell type mixtures in spatial transcriptomics. Nature Biotechnology, 40(4), 517–526.

[17] Andersson, A., Bergenstråhle, J., Asp, M., Bergenstråhle, L., Jurek, A., Fernández Navarro, J., & Lundeberg, J. (2020). Single-cell and spatial transcriptomics enables probabilistic inference of cell type topography. Communications biology, 3(1), 1–8.

[18] Kleshchevnikov, V., Shmatko, A., Dann, E., Aivazidis, A., King, H. W., Li, T., Elmentaite, R., Lomakin, A., Kedlian, V., Gayoso, A., et al. (2022). Cell2location maps fine-grained cell types in spatial transcriptomics. Nature biotechnology, 40(5), 661–671.

[19] Ma, Y., & Zhou, X. (2022). Spatially informed cell-type deconvolution for spatial transcriptomics. Nature Biotechnology, 1–11.

[20] Tasic, B., Yao, Z., Graybuck, L. T., Smith, K. A., Nguyen, T. N., Bertagnolli, D., Goldy, J., Garren, E., Economo, M. N., Viswanathan, S., et al. (2018). Shared and distinct transcriptomic cell types across neocortical areas. Nature, 563(7729), 72–78.

[21] Maynard, K. R., Collado-Torres, L., Weber, L. M., Uytingco, C., Barry, B. K., Williams, S. R., Catallini, J. L., Tran, M. N., Besich, Z., Tippani, M., et al. (2021). Transcriptome-scale spatial gene expression in the human dorsolateral prefrontal cortex. Nature neuroscience, 24(3), 425–436.

[22] Pardo, B., Spangler, A., Weber, L. M., Page, S. C., Hicks, S. C., Jaffe, A. E., Martinowich, K., Maynard, K. R., & Collado-Torres, L. (2022). Spatiallibd: An r/bioconductor package to visualize spatially-resolved transcriptomics data. BMC genomics, 23(1), 434.

[23] Huuki-Myers, L. A., Spangler, A., Eagles, N. J., Montgomery, K. D., Kwon, S. H., Guo, B., Grant-Peters, M., Divecha, H. R., Tippani, M., Sriworarat, C., et al. (2023). Integrated single cell and unsupervised spatial transcriptomic analysis defines molecular anatomy of the human dorsolateral prefrontal cortex. BioRxiv, 2023–02.

[24] Wu, R., Guo, W., Qiu, X., Wang, S., Sui, C., Lian, Q., Wu, J., Shan, Y., Yang, Z., Yang, S., et al. (2021). Comprehensive analysis of spatial architecture in primary liver cancer. Science Advances, 7(51), eabg3750.

[25] Jiang, Z., Wu, Y., Miao, Y., Deng, K., Yang, F., Xu, S., Wang, Y., You, R., Zhang, L., Fan, Y., et al. (2023). Hccdb v2. 0: Decompose the expression variations by single-cell rna-seq and spatial transcriptomics in hcc. bioRxiv, 2023–06.

[26] Calderaro, J., Petitprez, F., Becht, E., Laurent, A., Hirsch, T. Z., Rousseau, B., Luciani, A., Amaddeo, G., Derman, J., Charpy, C., et al. (2019). Intra-tumoral tertiary lymphoid structures are associated with a low risk of early recurrence of hepatocellular carcinoma. Journal of hepatology, 70(1), 58–65.

[27] Kuang, D.-M., Zhao, Q., Peng, C., Xu, J., Zhang, J.-P., Wu, C., & Zheng, L. (2009). Activated monocytes in peritumoral stroma of hepatocellular carcinoma foster immune privilege and disease progression through pd-l1. Journal of Experimental Medicine, 206(6), 1327–1337.

[28] Kuang, D.-M., Peng, C., Zhao, Q., Wu, Y., Chen, M.-S., & Zheng, L. (2010). Activated monocytes in peritumoral stroma of hepatocellular carcinoma promote expansion of memory t helper 17 cells. Hepatology, 51(1), 154–164.

